# Generalized distribution interpolated between the exponential and power-law distributions and applied to pill bug (*Armadillidium vulgare*) walking data

**DOI:** 10.1101/2021.11.29.470497

**Authors:** Shuji Shinohara, Hiroshi Okamoto, Toru Moriyama, Yoshihiro Nakajima, Takaharu Shokaku, Akika Utsumi, Ung-il Chung

**Affiliations:** School of Science and Engineering, Tokyo Denki University, Saitama, Japan; Department of Bioengineering, Graduate School of Engineering, The University of Tokyo, Tokyo, Japan; Faculty of Textile Science, Shinshu University, Ueda, Japan; Graduate School of Economics, Osaka City University, Osaka, Japan; Department of Network Design, Meiji University, Tokyo, Japan

**Keywords:** power-law distribution, exponential distribution, generalized distribution, gamma distribution, Weibull distribution, time dependence of step-length, pill bug, random walk

## Abstract

The Lévy walk, a type of random walk in which the frequency of linear step-lengths follows a power-law distribution, can be observed in the migratory behavior of organisms at various levels, from bacteria and T cells to humans. Compared to the Brownian walk, which is also a type of random walk (characterized by an exponential distribution of the frequency of occurrence of step-length), the Lévy walk is characterized by the occasional appearance of linear movements over very long distances. In this paper, we propose a general distribution that includes the power-law and exponential distributions as special cases. This distribution has two parameters: the first parameter represents the exponent, similar to the power-law and exponential distributions and the second is a shape parameter representing the shape of the distribution. By introducing this distribution, an intermediate distribution model can be interpolated between the power-law and exponential distributions. The shape parameter measures whether a distribution assimilates a power-law or exponential distribution. In this study, the proposed distribution was fitted to the frequency distribution of the step-length calculated from the walking data of pill bugs. The autocorrelation coefficients were also calculated from the time-series data of the step-length, and the relationship between the shape parameter and time dependency was investigated. The results showed a significant negative correlation between the two (*r*=−0.61, n=30, t=4.04, and p=0.00037). This means that pill bugs with gait patterns closer to Lévy than Brownian walks have a stronger time dependence with respect to step-length changes.

## 1. Introduction

Lévy walks are found in the migratory behavior of organisms at various levels, from bacteria and T cells to humans [1–6]. Lévy walks are a type of random walk in which the frequency of occurrence of a linear step-length *l* follows a power-law distribution *p* (*1*) □ *1* ^−*μ*^,1 < *μ* ≤ 3 . Compared to the Brownian walk, which is also a type of random walk (characterized by an exponential distribution *p* (*1*)□D *e*^−*λl*^ of the frequency of occurrence of step-length *l*), the Lévy walk is characterized by the occasional appearance of linear movements over very long distances.

Generally, Lévy walks with exponents close to two have been observed in the migratory behavior of organisms, and attention has been paid to why such patterns occur [7–11]. The Lévy flight foraging hypothesis (LFFH) [12][13] states that if food is sparse and randomly scattered and predators have no information (memory) about the food, Lévy walks, as a random search, will be the optimal foraging behavior and will be evolutionarily advantageous [14]. Until now, it has been considered that search efficiency is maximized for inverse square Lévy walks (also called Cauchy walks) with an exponent of two in the LFFH [15]. However, recent studies have indicated that a Cauchy walk does not necessarily have maximum search efficiency in spaces of two or more dimensions, as this is true only under special conditions [16]. In contrast, Guinard and Korman proved that an intermittent Cauchy walk is an optimal search strategy in finite two-dimensional domains when the goal is to rapidly find targets of arbitrary sizes [17]. Debates about natural conditions and search methods that make the Cauchy walk optimal are ongoing.

To determine whether the gait pattern is a Lévy walk or a Brownian walk, a comparison is made as to whether the frequency distribution of the linear step-length follows a power-law distribution or an exponential distribution [5,18-22]. In the comparison, the range of step-lengths to be analyzed and the parameters of the model that best fit the data in that range, that is, the exponents μ and λ, are first calculated. The maximum likelihood estimation method is generally used to estimate the parameters. Next, a comparison is made to determine which model fits better, the power-law distribution model, or the exponential distribution model. For comparison, Akaike information criteria weights (AICw), which consider the likelihood and number of parameters, are often used [21,22]. To verify whether the model fits the observed data, Clauset et al. [20] proposed goodness-of-fit tests based on the Kolmogorov–Smirnov (KS) statistic. In this way, judgments have been made as to whether the observed data follow a power-law distribution or an exponential distribution model, but many actual data cannot be classified as either [3].

In this paper, we propose a general distribution that includes the power-law and exponential distributions as special cases. This distribution has two parameters: the first parameter represents the exponent, similar to the power-law and exponential distributions and the second is a shape parameter representing the shape of the distribution. In this distribution, if the shape parameter is set to zero, it represents the power-law distribution, and if it is set to one, it represents an exponential distribution. By introducing this distribution, distributions that are intermediate between the power-law and exponential distributions can be modeled. In this study, the proposed distribution was fitted to the walking data of pill bugs collected by Shokaku et al. [3] as specific observation data. Differently expressed, we estimated the exponent and shape parameters that best fit the observed data.

It has been asserted that there is a time dependency in the time-series of the step-length in human mobility behavior. For example, Wang et al. [23], Rhee et al. [24], and Zhao et al. [25] demonstrated that the temporal variation of step-length is autocorrelated, which means that there is a trend in the time variation of step-length, such that short (long) steps are followed by short (long) steps. In addition, it has been emphasized that the time dependence of step-length is related to the frequency distribution of the step-length following a power-law distribution [23].

In this study, to investigate the time dependence of the walking data of the pill bugs described above, we calculated the autocorrelation coefficient between the time-series *l*_*t*_ of the step-length and the time-series *l*_*t* +*τ*_ with time lag τ. When τ=0, the autocorrelation coefficient is one because the two time-series are identical. Conversely, when τ>0, any random walk such as the Lévy or Brownian walk is theoretically uncorrelated. We investigated the relationship between the autocorrelation coefficient calculated from the time-series data of the linear step-length and the shape parameter of the frequency distribution of the linear step-length. The result showed a significant negative correlation between the two (*r*=−0.61, n=30, t=4.04, and p=0.00037). This means that pill bugs with gait patterns closer to Lévy than Brownian walks have a stronger time dependence with respect to step-length changes.

## 2. Methods

### 2.1. A distribution interpolated between the exponential and power-law distributions

In this section, we propose a general distribution that includes exponential and power-law distributions as special cases. First, let us consider the process of elongating the persistence length. This length can be a spatial distance or a time interval. For example, consider a process in which an organism continues to move in a straight line and reaches a distance of *l*, and then moves in the same straight line by a *dl*, for the total straight-line distance to increase to *l* + *dl*. As a discrete example, consider the process of tossing a coin and getting “heads” *l* times, then getting “heads” again, and extending the period of consecutive “heads” to *l*+1 times.

Suppose that the probability of length *l* occurring is represented by *p*(*l*). Because *l* is a length, let *l*≥0. Because *p*(*l*) is a probability distribution, it satisfies 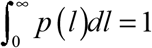. Let us also denote by *P*(*l*) the complementary cumulative frequency distribution (CCDF) in which lengths greater than or equal to *l* occur. That is,

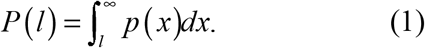

Based on this definition,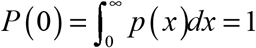 is valid. Similarly, from the definition, the following relationship holds between *p*(*l*) and *P*(*l*).

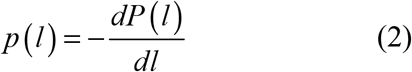

We denote *f*(*l*) the probability that the length *l* will extend to *l* + *dl*. *f*(*l*) can be expressed by the following equation:

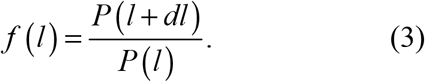

In contrast, the probability that length *l* is reached and ends at *l* is denoted by *g*(*l*). *g*(*l*) can be expressed using *p*(*l*) and *P*(*l*) as follows:

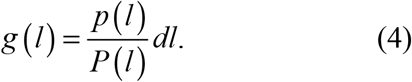

Here, an interval of length *l* can either extend further from *l* or end at *l*, therefore, *f*(*l*)+*g*(*l*)=1 holds. To consider the elongation process in the discrete case, that is, when *dl*=1, let us consider a situation in which a coin is tossed repeatedly. We assume a situation in which the probability of “heads” in a coin toss is not constant, but varies depending on the number of times the same side appears consecutively. For example, after three consecutive “heads,” the probability that the next is also “heads” is represented by *f*(3).Conversely, the probability that the next is “tails” is expressed as *g*(3)=1−*f*(3).

The CCDF that the same side will continue for *l*+1 or more consecutively is expressed as follows:

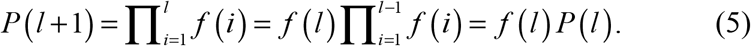

The probability that the same side continues consecutively for *l* times, that is, the probability of length *l* occurring, is expressed as follows:

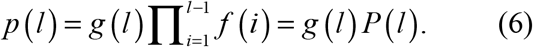

Now, let us consider Brownian and Lévy walks. For the Brownian walk, the probability of a step-length *l* occurring and its CCDF is represented by the following exponential distribution with an exponent *β*.

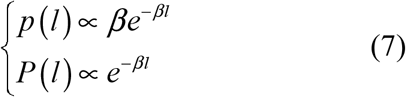

Conversely, for the Lévy walk, the probability of the occurrence of a step-length *l* and the CCDF are expressed by the following power-law distributions with the exponent *β*+1 and *β*, respectively:

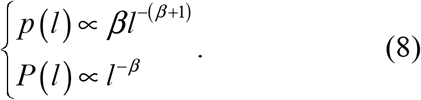

If the change rate (failure rate) at length *l* is defined as 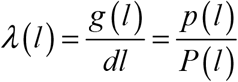, the change rates of the Brownian walk and the Lévy walk can be expressed as follows:

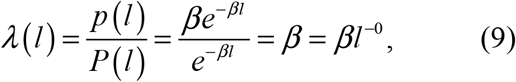

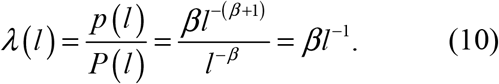

We now define the generalized rate of change to include the Brownian and Lévy walk as a special case, as follows:

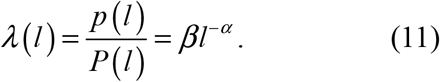

The case of α=0 corresponds to the change rate of the Brownian walk, and the case of α=1 corresponds to the change rate of the Lévy walk.

Equation (11) can be written as 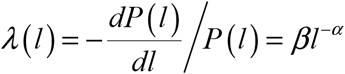, and solving it for *P* (*l*) yields the following solution:

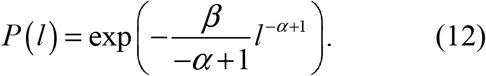

If we put *m* = −*α* +1, then *P* (*l*) is expressed as follows:

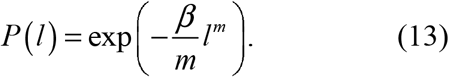

Equation (11) can be rewritten as follows:

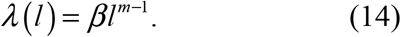

The probability *p*(*l*) of step-length *l* occurring can be written as follows, using Equation (2) and (13).

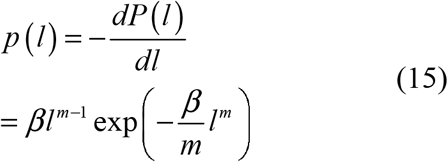

Let us consider the meanings of *m* and *β*. As can be seen from Equation (14), the change rate becomes smaller as the value of *l* increases, since *m*−1<0 holds in the range of *m*<1. Moreover, the smaller the value of *m*, the stronger the tendency. This may be viewed as a process of gradual acclimatization to going straight ahead. When *m*>1, the change rate increases as *l* increases. This may be viewed as a change due to fatigue or boredom from continuing straight ahead. When *m*=1, the change rate is a constant value *β*, regardless of the value of *l*. In other words, this change was accidental. Thus, *m* is a parameter that controls how the length of *l*, or the past history of how long the same condition has lasted is considered.

Conversely, for *β*, if *l* and *m* are fixed, the larger *β* becomes, the higher the change rate becomes. Otherwise expressed, *β* is a parameter that controls the magnitude of the change rate. For *m* = 1, Equation (15) is expressed as follows:

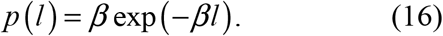

Put differently, it represents an exponential distribution of the exponent *β*. Conversely, when *m* = 0, Equation (15) cannot be defined because it involves division by zero. However, when *m* is sufficiently close to zero, this distribution can be approximated using the Maclaurin expansion, *l*^*m*^ ≈ 1+ *m* log *l* as follows:

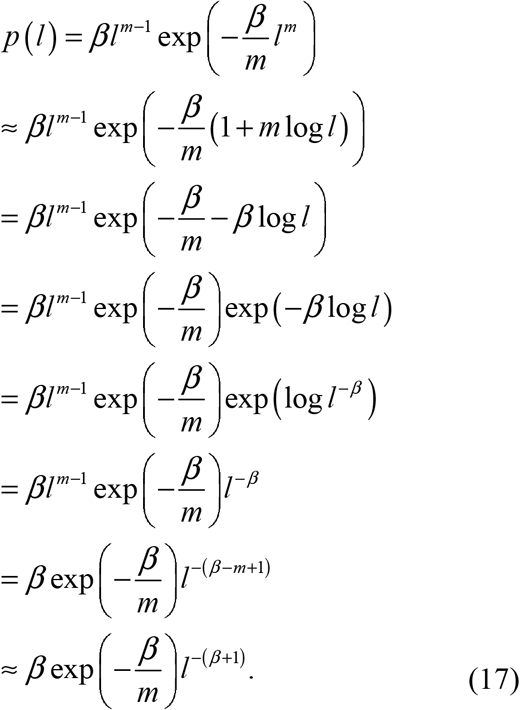

If we set 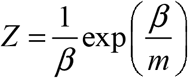 as the normalization constant, Equation (17) can be rewritten as follows:

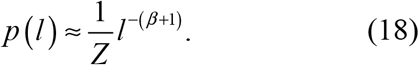

Otherwise expressed, Equation (15) represents the power-law distribution of the exponent *β*+1 as an approximation. Thus, Equation (15) can be said to be a distribution that includes not only the exponential distribution but also the power-law distribution as a special case, albeit as an approximation. Therefore, in this study, for convenience, the distribution of Equation (15) is named the generalized distribution (GE). Furthermore, parameter *m* is called the shape parameter in the sense that it represents the shape of the distribution.

### 2.2 Relationship between GE and Weibull distribution

This section describes the relationship between GE and the Weibull distribution (WB). The generalized gamma distribution is defined using three parameters—*m, s*, and η—as follows:

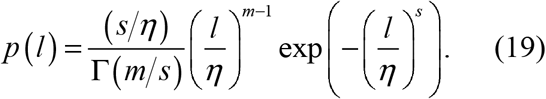

where Г (·) denotes the gamma function; for *s* = *m*, the generalized gamma distribution represents the Weibull distribution (WB) [26] as follows:

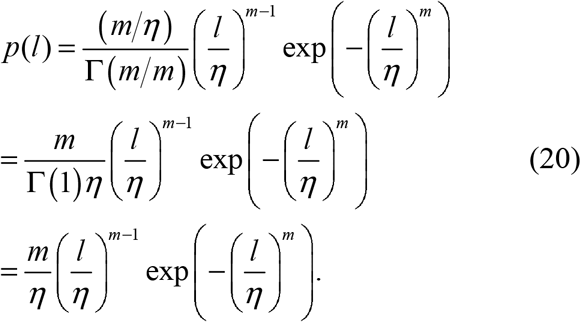

Here, η is the scale parameter. *m* is a parameter that determines the shape of the distribution and is called the shape parameter.

For *m*=1, the WB represents the exponential distribution of the scale parameter η.

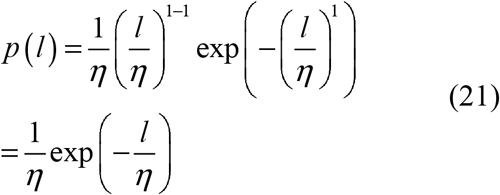

The WB thus includes the exponential distribution as special case.

To compare the WB with the GE here, a scale parameter η is introduced to the GE presented in Equation (15) and extended to three variables as follows:

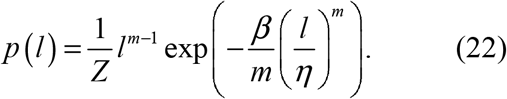

where *Z* is assumed to represent the normalization constant. Note that if η = 1, Equation (22) coincides with Equation (15), i.e. GE.

Equation (20) can be transformed with *Z* as the normalization constant as follows:

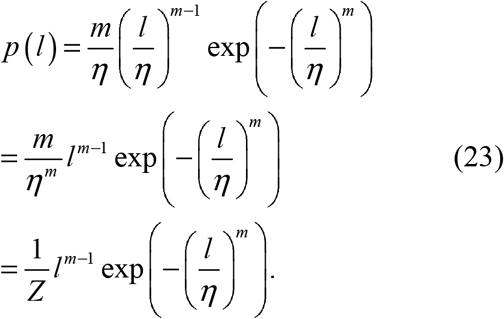

Comparing Equations (22) and (23), the WB corresponds to the case with constraint *β* = *m* in Equation (22). In the case of *m* → 0, Equation (22) can be transformed to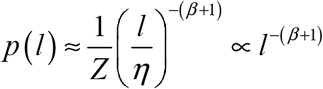, which can be approximated as a power-law distribution with exponent *β*+1, as shown in Equation (15). For *m* → 0, the WB approximates a power-law distribution with exponent 1 because of constraint *β* = *m*.

### 2.3 GE and entropy maximization

In the following section, we analyze GE from the perspective of entropy maximization [27]. First, the entropy of the probability distribution is defined as follows:

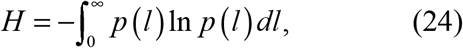

Where *p* (*l*) is a probability distribution and thus satisfies the following constraints:

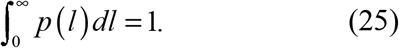

Second, we define the generalized mean of the probability distribution as follows:

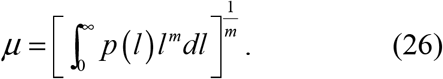

For *m* = 1 and *m* = −1, Equation (26) represents the arithmetic and harmonic means, respectively. In the case of *m* = 0, Equation (26) cannot be defined because it includes a division by zero. However, using the McLaurin expansion for *m* → 0, Equation (26) can be approximately transformed as follows:

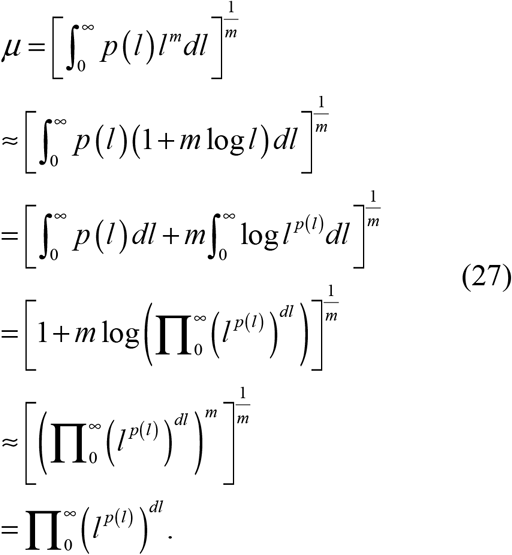

That is, for *m* → 0, Equation (26) represents an approximate geometric mean.

We find the probability distribution that maximizes the entropy under a constant mean constraint. In other words, we find the probability distribution that maximizes Equation (24) under the constraints of Equations (25) and (26). To do this, we find *p* (*l*) by maximizing the following *L*, using Lagrange’s undetermined multiplier method:

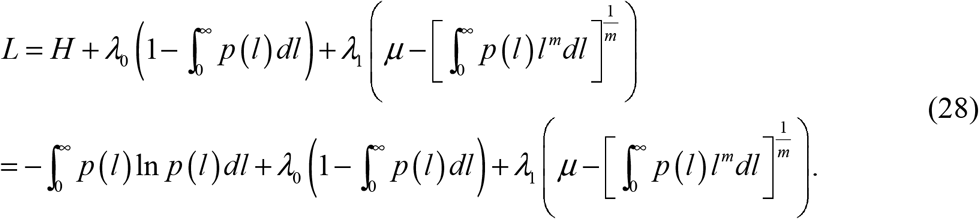

The partial derivative of *L* can be calculated as follows:

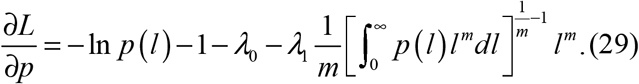

Since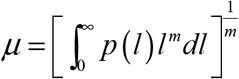, Equation (29) is transformed as follows:

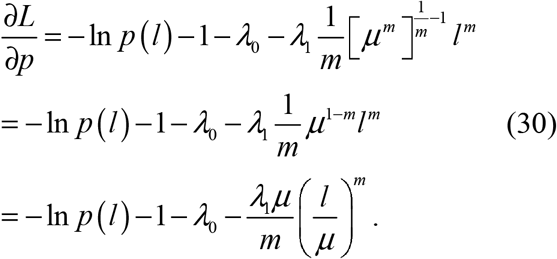

Solving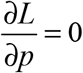, *p* (*l*) is calculated as follows:

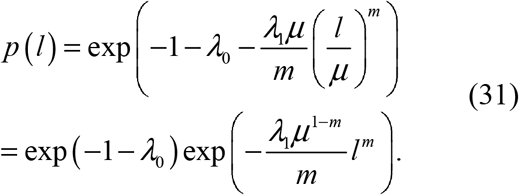

We let *λ μ*^1−*m*^ = *β*and 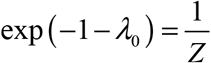, then

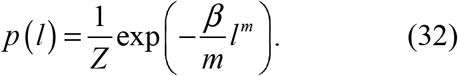

Equation (32) is a distribution that maximizes entropy when constrained such that the generalized mean is constant. It can be seen that this equation takes the same form as the CCDF of GE expressed in Equation (13). When *m* = 1, that is, the arithmetic mean is constant, Equation (32) becomes 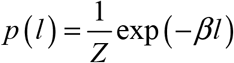, which is an exponential distribution. When *m* → 0, that is, the geometric mean is constant, Equation (32) becomes 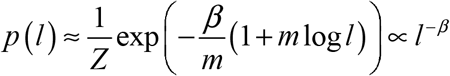, which is a power-law distribution with exponent *β*.

### 2.4. Walking data of pill bugs

We applied the method described above to the pill bug’s gait data. It is known that pill bugs have a habit termed turn alternation, following which they turn to the right (left), left (right), and so on [28]. The mechanism underlying turn alternation is assumed to be based primarily on proprioceptive information from the previous turn and arises from bilaterally asymmetrical leg movements that occur when turning [29]. During one turn, the outer-side legs travel further than the inner ones. After completing the turn, the relatively rested inner-side legs exert more influence on subsequent movements than the outer-side ones and bias the animal to turn in the opposite direction at the next step.

By alternating turns, pill bugs can maintain a straight course to avoid an obstacle. Moving in a straight course is considered the most adaptive strategy when precise information about environmental resources or hazards is absent [30]. However, when pill bugs were examined in successive T-mazes, they sometimes turned in the same direction as they had at the previous junction (turn repetition). For example, in an experiment on 12 pill bugs using 200 successive T-mazes (for approximately 30 min), three individuals maintained a high rate of turn alternation, four a low rate, and the remaining five spontaneously increased and decreased the rate [31]. Why some pill bugs did not maintain a high rate of turn alternation, that is, generate turn repetition at a rate other than low, is still unclear.

Shokaku et al. developed an automatic turntable-type multiple T-maze device to observe the appearance of turn alternation and turn repetition in pill bugs over a long period and to investigate the effects of these turns on gait patterns [3]. This is a virtually infinite T-maze that uses a turntable. The pill bug turns to the left or right at a T-junction, goes straight ahead, and then crosses another T-junction, and so on. Using this device, Shokaku et al. observed 33 pill bugs for more than 6 h each. An example of a walking pattern in the T-maze is shown in Figure 1.

**Figure 1.**
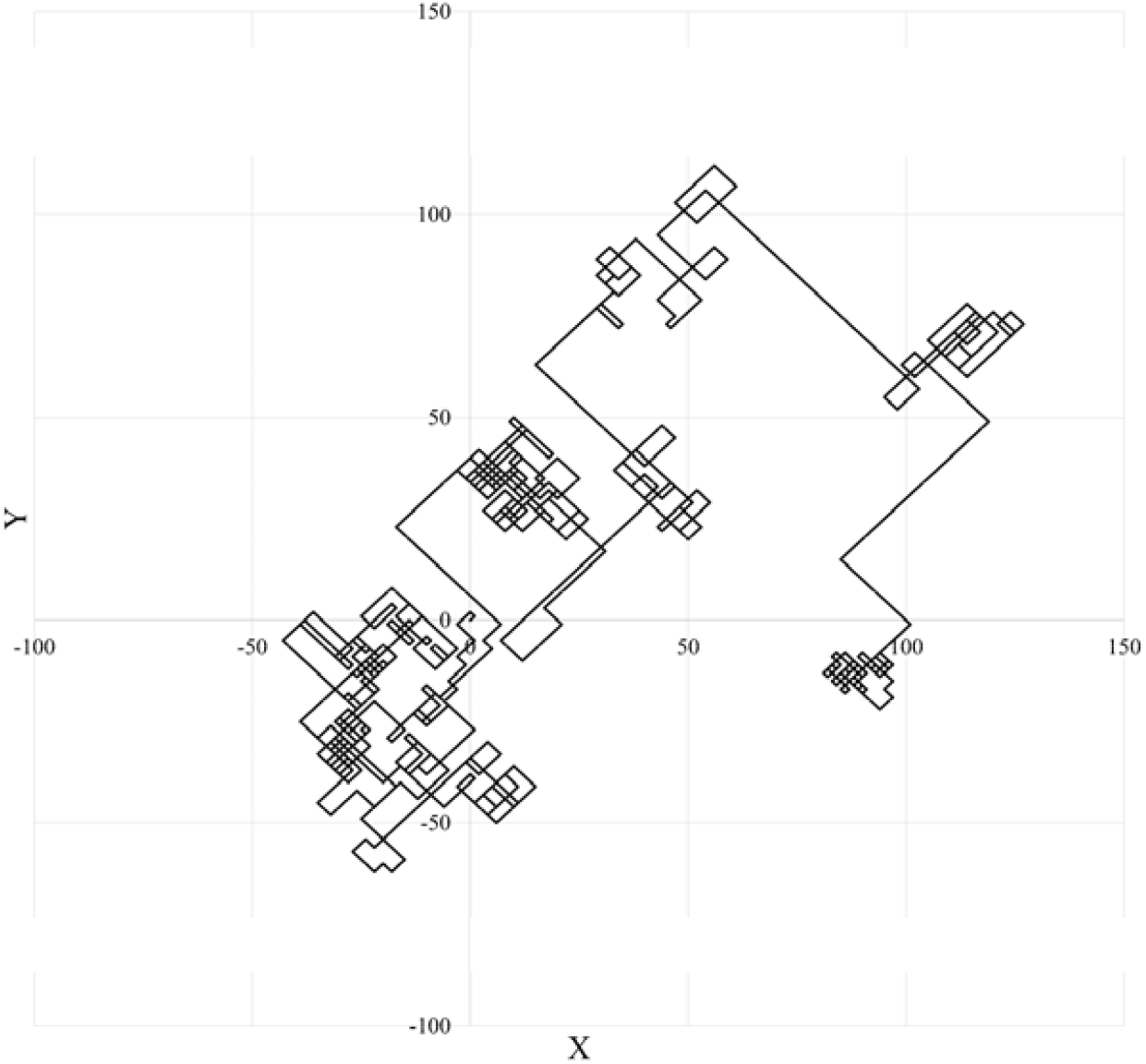
Trajectory of an individual’s gait.

When the turns are repeated regularly, alternating left and right, the pill bug is considered to move straight. Conversely, if the same turn is repeated, such as right and right, it is considered to have changed direction.

In this classification of gait patterns, the pill bug is considered to decide whether to continue or abort straight-ahead movement each time it encounters a T-intersection. The straight-line distance *l* was calculated using the method shown in Figure 2. Using this method, time-series data of the straight-line distance *l* for each individual were obtained.

**Figure 2.**
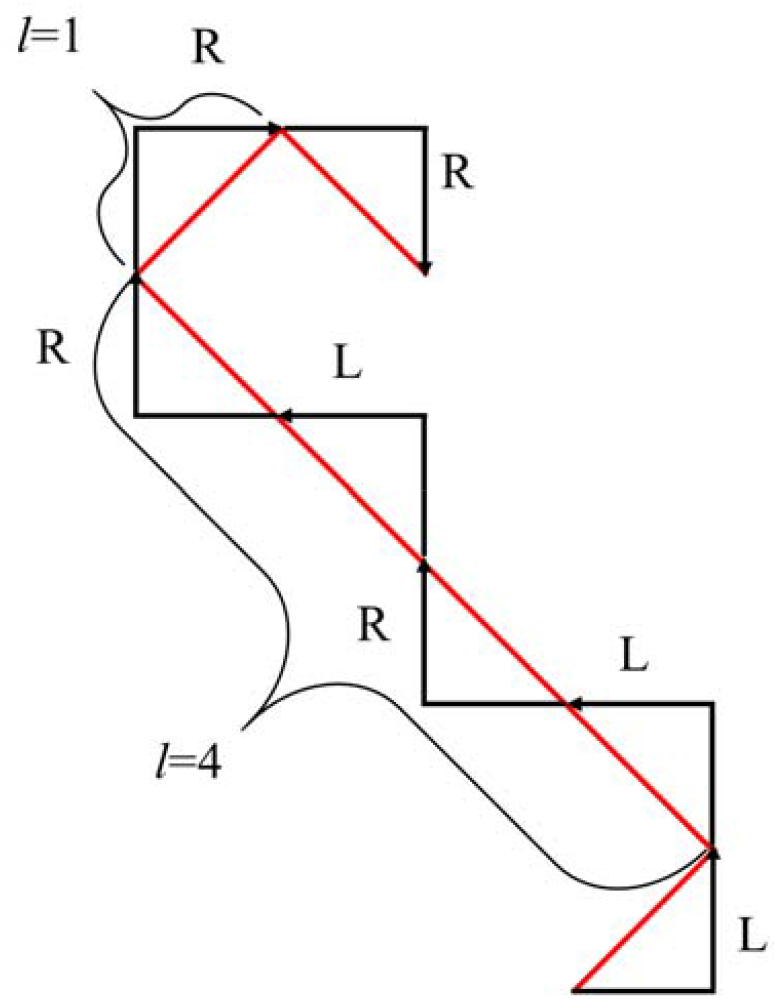
Sample calculation of step-length *l*. The black polygonal line with the arrow represents a turn; L represents a left turn, R a right turn. The red line represents an approximate linear movement. The L–R– L–R pattern shown in this figure represents linear movement with a step-length of 4 (*l* = 4).

### 2.5 Fitting GE to the walking data of pill bugs

We describe a method to fit the frequency distribution of the step-length observed in pill bugs to the generalized distribution (GE). The method was based on references [5,18-21].

The first objective of this section is to find the minimum 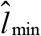 and maximum 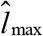 of the observed data that should be fitted to the model. The second objective is to find the parameters of the model that best fit the data in the range 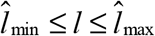.

Suppose we are given *N* observed data *D* = {*l*_1_,*l*_2_,?,*l*_*N*_ } in the range of *l*_min_ ≤ *l* ≤ *l*_max_ . The model for the data must be satisfied 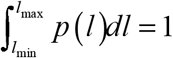. For this reason, we redefine the GE model as follows:

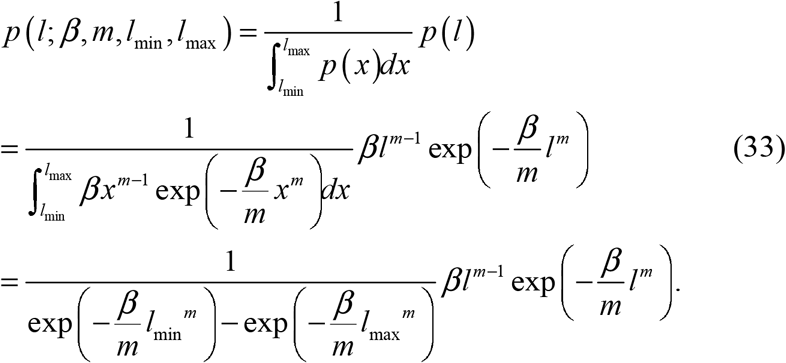

When the observed data are discrete, such as natural numbers, they are defined as follows:

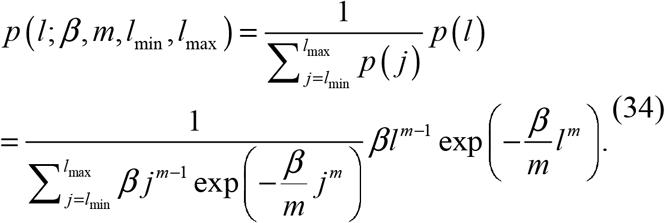

In addition, because the CCDF of *p* (*l*; *β, m, l*_min_, *l*_max_) must satisfy *P* (*l*_min_) = 1, we redefine it as follows:

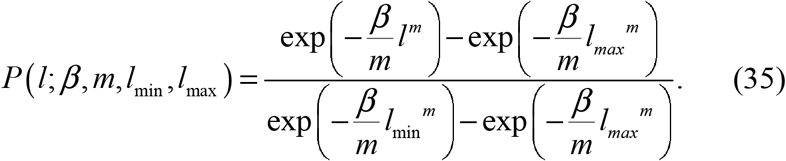

When the observed data is discrete, the CCDF is defined as follows:

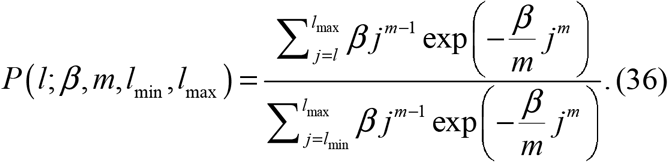

The log-likelihood of the observed data *D* was calculated using Equation (33) as follows:

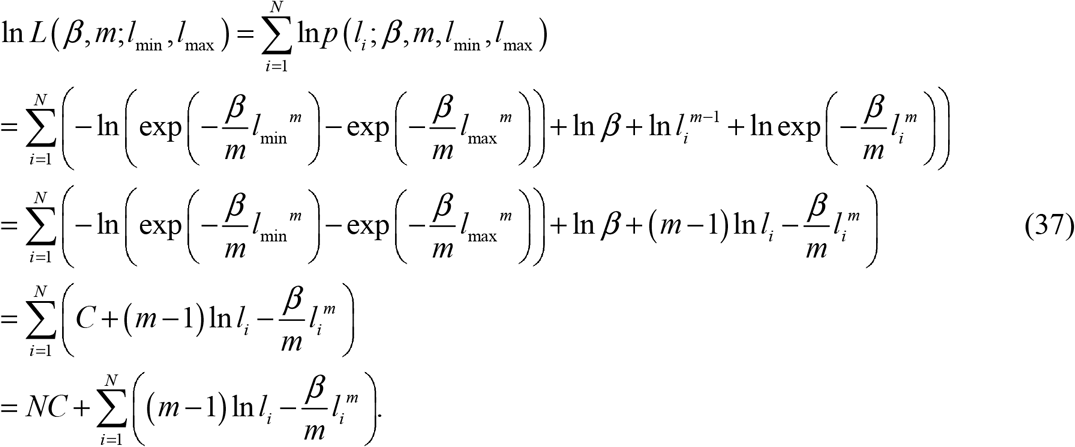

Here, 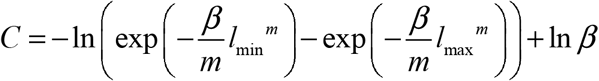.The log-likelihood when the observed data are discrete can be replaced by 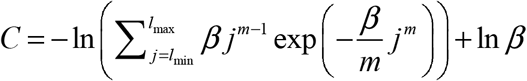.

The model parameters that best fit the observed data are 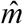 and 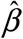, which maximize the log-likelihood. In this paper, *m* varies from −1.5 to 1.5 and *β* from 0.01 to 4.0 in increments of 0.01, and the parameters 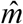 and 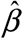 for which Equation (37) is maximized are obtained numerically. When *m* = 0, Equation (37) cannot be defined; therefore, we treat GE as the power-law distribution.

To evaluate the model’s goodness-of-fit to the observed data, we used the Kolmogorov–Smirnov statistic *D* (*l*_min_, *l*_max_), which represents the distance between the CCDF, *S* (*l*;*l*_min_, *l*_max_), calculated from the observed data *D* and the theoretical CCDF expressed in Equation (36).

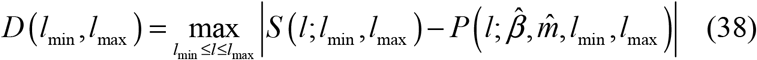

In this paper, let 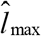 be the maximum *l*_max_ of the observed data. If 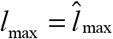, then 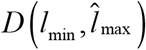 can be considered as a function of 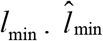, which minimizes 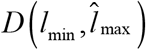, is numerically calculated from within the observed data. That is,

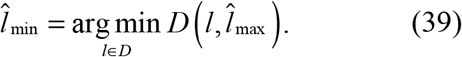

As shown above,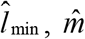, and 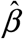 can be obtained numerically using Equation (37) and Equation (39).

### 2.6. Autocorrelation coefficient

In this study, we investigate the relationship between the shape of the frequency distribution of step-length and its time dependence. To quantify the former, the shape parameter *m* of the GE described above is used. To quantify the latter, the autocorrelation coefficient and average correlation coefficient of the step-length time-series are used.

Suppose we are given *N* time-series data (*l*_1_,*l*_2_,L, *l*_*N*_) . In this case, the autocorrelation coefficient *r*(τ) with time lag τ is expressed as follows:

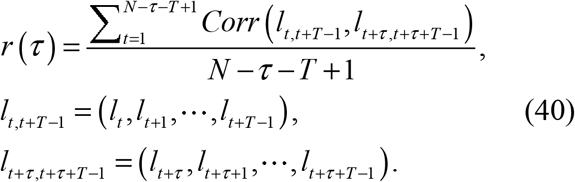

where *Corr* (*x, y*)is the correlation coefficient between the time-series data *x* and *y*. In this study, *T* = 200was set. When τ=0, we take the correlation coefficient between the same time-series data *r*(0)=1. We calculated the autocorrelation coefficients of the time-series data of linear step-length to investigate the time dependence of the pill bugs’ walking data. In this study, the autocorrelation coefficient was obtained in the range of τ = 1–50. The autocorrelation coefficients for every individual are averaged from *r*(1) to *r*(50) and are expressed as 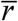 .

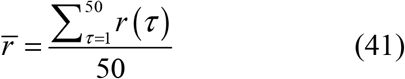

Thereafter, 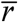 is referred to as the average autocorrelation coefficient.

### 2.7. Software used for analyses

The programs for the parameter estimation and autocorrelation coefficient calculation described above were developed using C++. The compiler was MinGW 8.1.0 64-bit for C++ [32]. The operating system used was Windows 10.

## 3. Results

Figure 3 shows examples of the GE. Figures 3(a) and 3(b) show the cases of *m*=1.0 and *m*=0.01, respectively. Figure 3(a) shows a single logarithmic graph with the vertical axis on a logarithmic scale and Figure 3(b) shows a double logarithmic graph with the vertical and horizontal axes on a logarithmic scale. In these figures, the GEs for the cases *β* = 1.0 and 2.0 are shown. In Figure 3(a), the exponential approximation curves are shown, and in Figure 3(b), the power approximation curves are shown. From the figures, we can see that the GE for *m* = 1.0, can be approximated by an exponential distribution with an exponent *β*. Conversely, the GE for *m* = 0.01 can be approximated by the power-law distribution with exponent *β*+1. Figure 3(c) shows the cases of *m*=0.5 and *m*=−0.5 for *β* = 1.0. This figure shows a double logarithmic graph with both axes on a logarithmic scale. As can be seen from the figure, GE shows an upward convex curve when *m*>0 and a downward convex curve when *m*<0.

**Figure 3.**
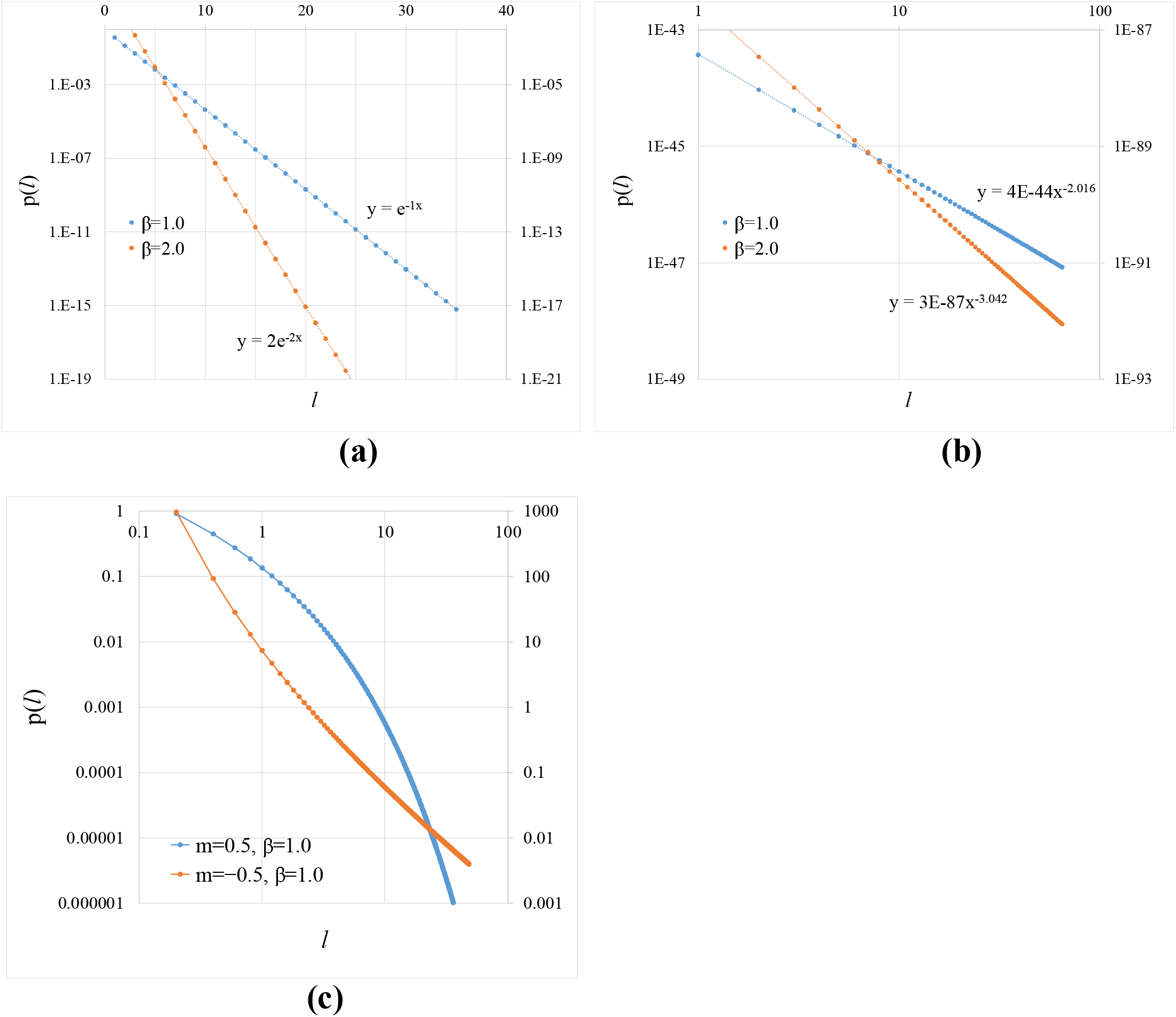
Examples of GE. (a) For *m* = 1.0, the GEs are shown for *β* = 1.0 and 2.0. The vertical axes are shown logarithmically. The left and right vertical axes represent the cases *β* = 1.0 and 2.0, respectively. The exponential approximation curves are also shown. (b) For *m* = 0.01, the GEs for *β* =1.0 and 2.0 are shown. The left and right vertical axes represent the cases *β* = 1.0 and 2.0, respectively. The vertical and horizontal axes are shown logarithmically. The power approximation curves are also shown. (c) For *β* = 1.0, the GEs for *m* = 0.5 and −0.5 are shown. The left and right vertical axes represent the cases *m* = 0.5 and −0.5, respectively. The vertical and horizontal axes are shown logarithmically.

Figure 4 shows examples of walking data for three individual pill bugs. Figure 4(a) shows the time-series of the step-length of subject 32 in the experiment of Shokaku et al [3]. Figure 4(b) shows the CCDF of the step-length. Figures 4(c) and 4(d) show the data for subject 16, and Figures 4(e) and 4(f) show the data for subject 34. Figures 4(b), (d), and (f) also show the results of fitting the CCDF of the GE model to the observed data. Figures 4(b), 4(d), and 4(f) are double logarithmic graphs with both axes displayed in logarithmic form. Note that in these figures, 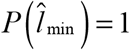 is based on the definition.

**Figure 4.**
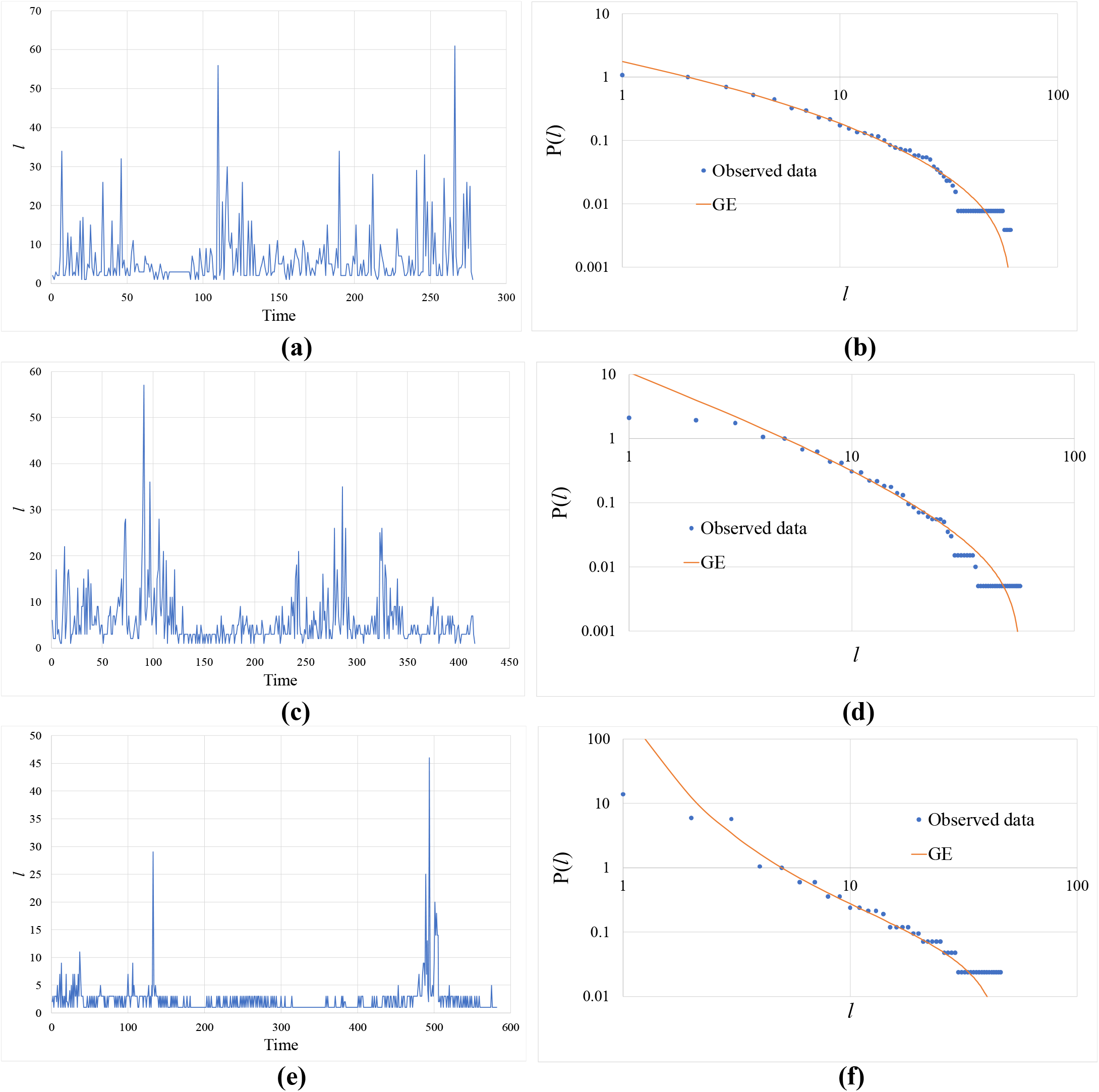
Step-length data for three individuals. (a) Step-length time-series for subject 32. (b) CCDF of subject 32. 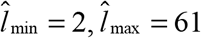, *m* = 0.32, *β* = 0.58 . (c) Step-length time-series for subject 16. (d) CCDF of subject 16. 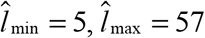, *m* = 0.20, *β* = 1.05 . (e) Step-length time-series for individual of subject 34. (f) CCDF of subject 34.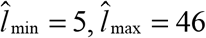, *m* = −0.56, *β* = 4.0

Figure 5 shows the correlograms for the three individual pill bugs shown in Figure 4. In this figure, the horizontal axis represents time lag τ. The vertical axis represents the autocorrelation coefficient *r*(τ) for τ.

**Figure 5.**
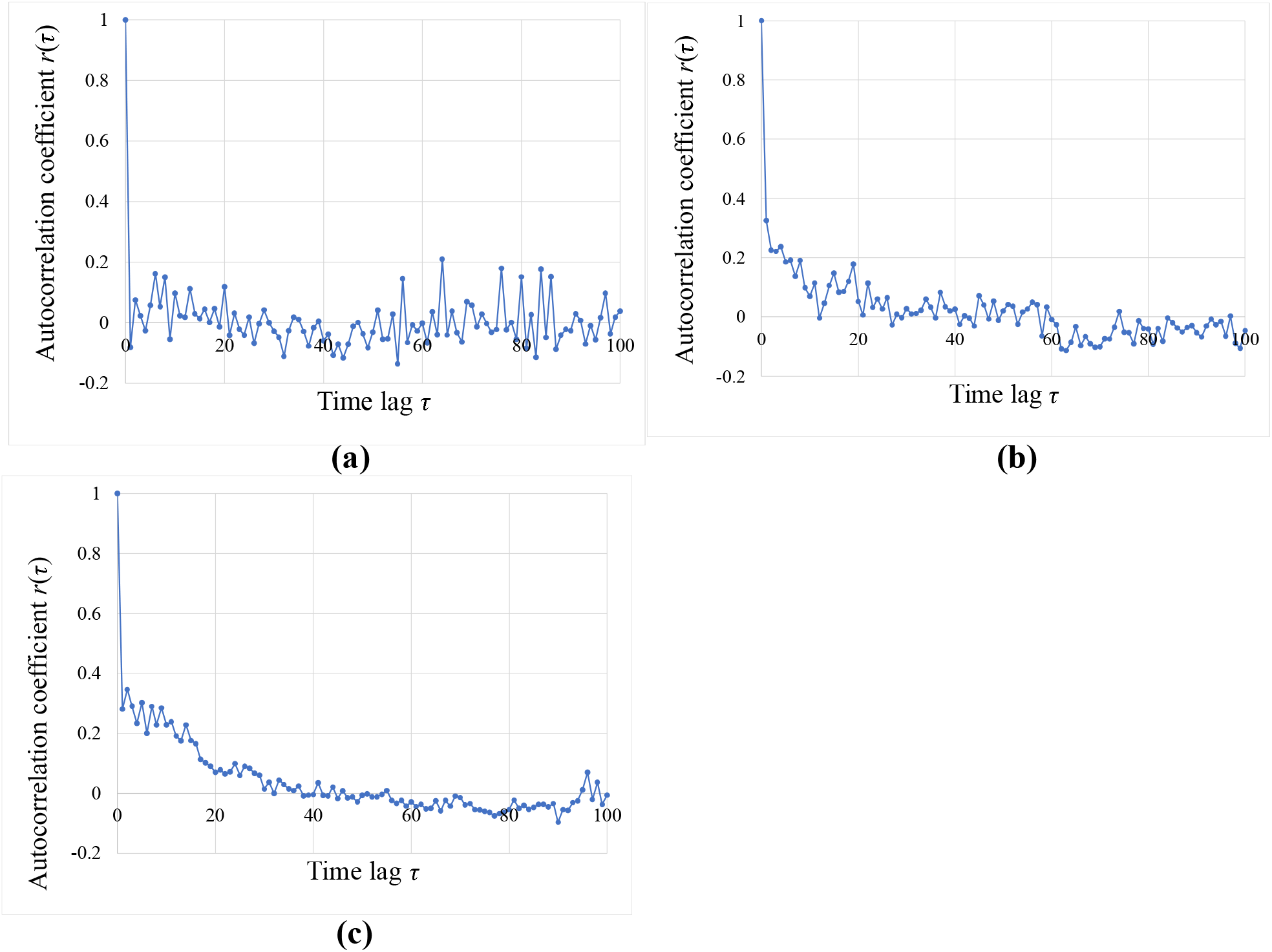
Autocorrelation of step-length time-series for three individuals. (a) Autocorrelation for subject 32. The average autocorrelation coefficient is 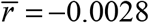. (b) Autocorrelation for subject 16. The average autocorrelation coefficient is 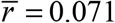. (c) Autocorrelation for subject 34. The average autocorrelation coefficient is 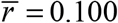.

Figure 6 shows the relationship between the shape parameter *m* when the GE fits the frequency distribution of each individual’s step-length and the time dependence of the time-series data of step-length. The autocorrelation coefficients were calculated from 1 to 50 for the time lag, and only individuals with more than 250 time-series data were included in the analysis. A total of 30 individuals were analyzed. The correlation coefficient between the shape parameters and average autocorrelation coefficients was −0.61 (*n*=30, *t*=4.04, and *p*=0.00037). In other words, the smaller the value of *m*, the stronger the time dependence of the time-series data of the step-length becomes.

**Figure 6.**
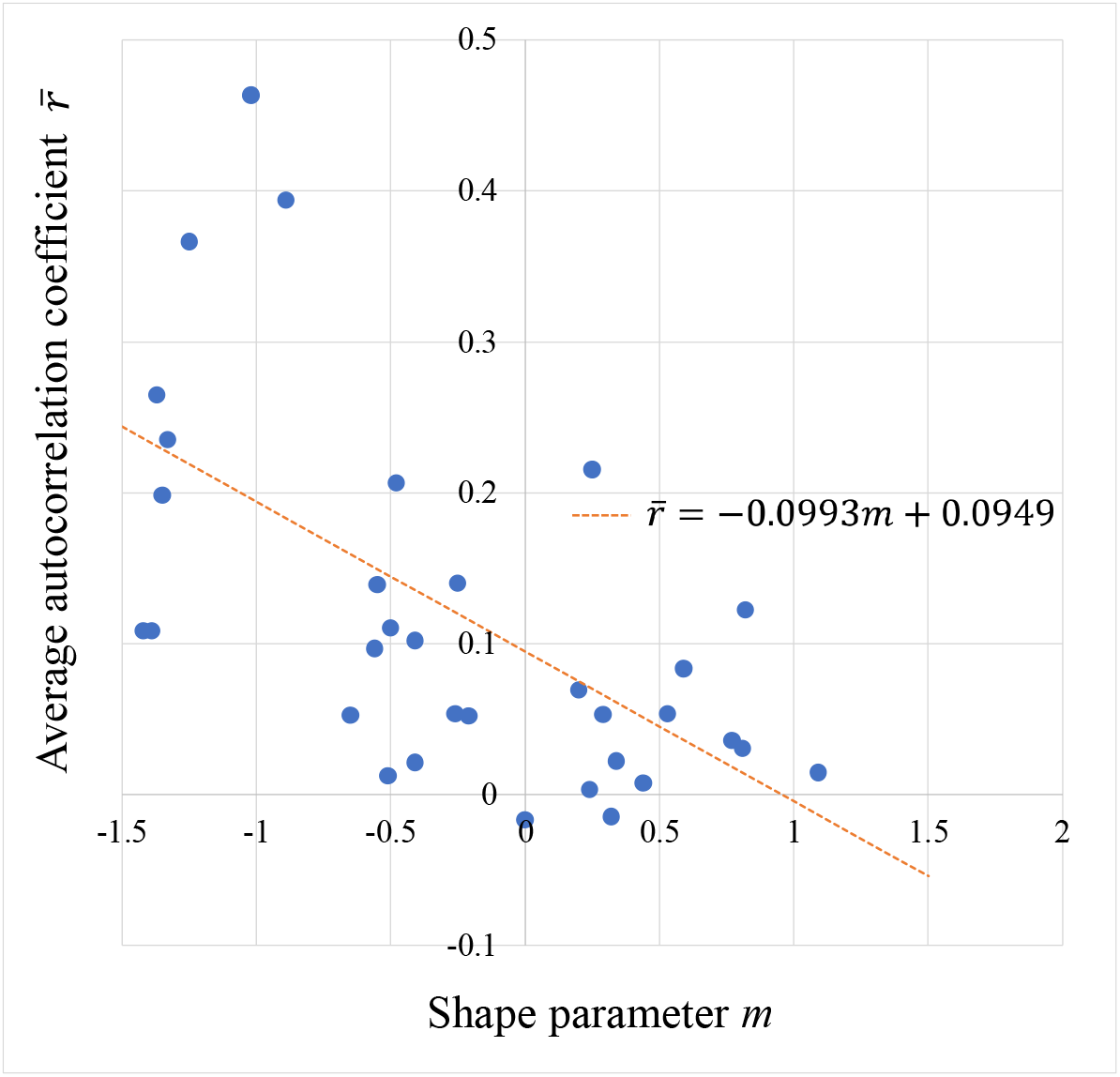
Relationship between shape parameters and average autocorrelation coefficients. The figure also includes a regression line.

Figure 7 shows the relationship between the shape parameter *m* and exponent *β* when the GE fits the frequency distribution of each individual’s step-length. The figure also shows the straight line *β*=−*m*+1. As can be seen from the figure, the data for some individuals line up near this straight line. In other words, the frequency distribution of these individuals can be approximated by a distribution with one parameter represented by 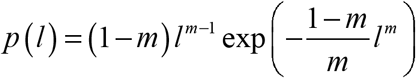. As shown in Equation (18), in the case of *m* → 0, GE can be approximated by a power-law distribution with exponent *β*+1. Therefore, if the relation *β*=−*m*+1 holds, then in the case of *m* → 0, it can be approximated by the power-law distribution with exponent two.

**Figure 7.**
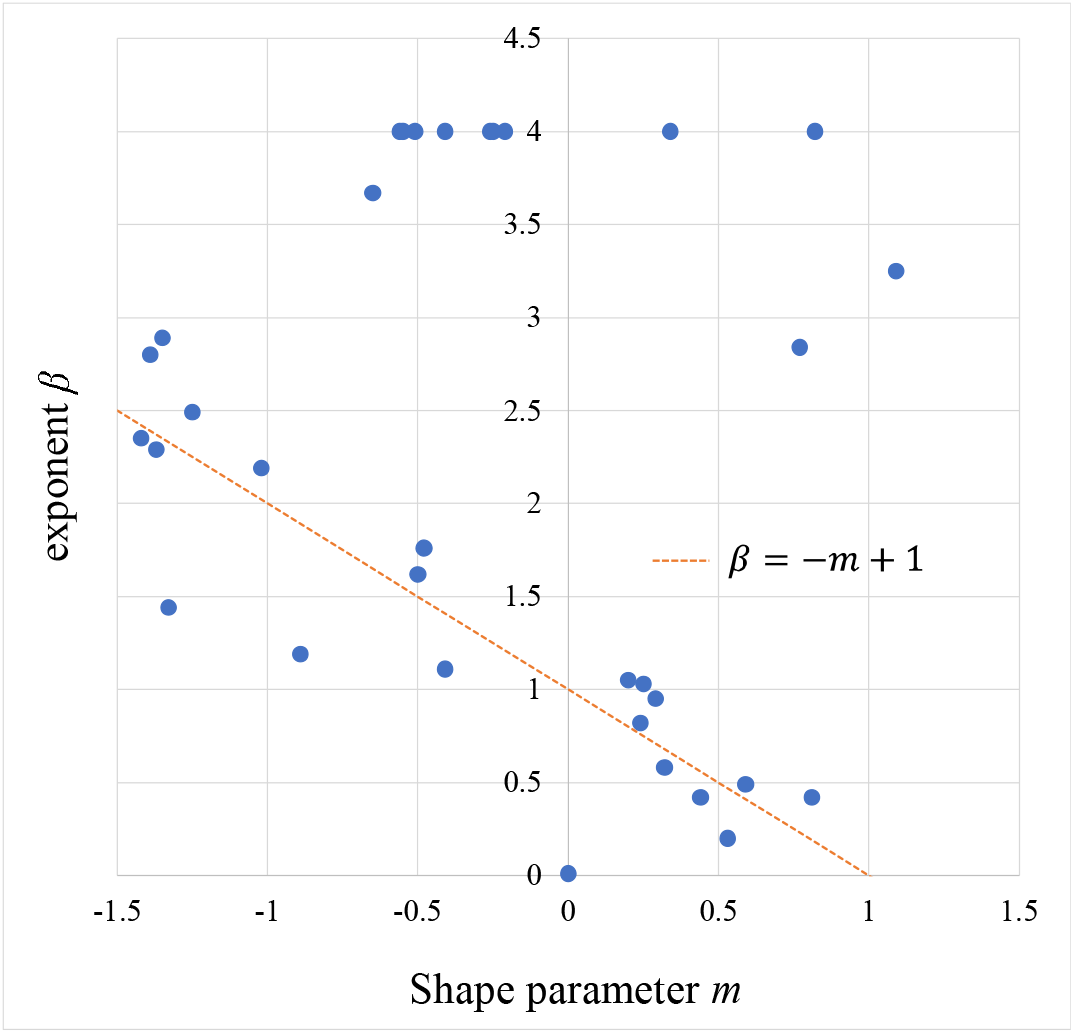
Relationship between shape parameters and exponents. The figure also includes the straight line *β*=−*m*+1.

## 4. Discussion

This study proposed a generalized distribution that includes exponential and power-law distributions as special cases. The proposed GE was applied to the walking data of pill bugs. A significant negative correlation was observed between the shape parameter *m* when the GE fitted the frequency distribution of each individual’s step-length and the time dependence of the time-series data of the step-length, as shown in Figure 6. However, this correlation is nontrivial. For example, if we randomly shuffle the order of the time-series data, as shown in Figure 4 (e), the time dependence disappears, but the value of the shape parameter is unchanged because the probability distribution, as shown in Figure 4 (f), is not affected by the shuffling.

Ross et al. [33] demonstrated that in human hunting behavior, the mode of exploration changes depending on encounters with prey. In particular, they indicated that in response to encounters, hunters more tortuously search areas of higher prey density and spend more of their search time in such areas; however, they adopt more efficient unidirectional, inter-patch movements after failing to encounter prey soon enough. This type of search behavior is called an area-restricted search (ARS) [33,34]. In ARS, searches with short travel distances within patches are combined with searches with long travel distances between patches, so there may be a tendency for relatively short and long straight distances to appear more frequently than intermediate straight distances, that is, distributions with small shape parameters may be more likely to appear. In addition, the time dependence of the step-length may appear because the search within a patch continues for a while after encountering the food, or conversely, the search between patches continues for a while when the food is not found. It may be that the power-law distribution and time dependence tend to appear simultaneously when the search between different levels is combined hierarchically, such as intra-patch and inter-patch searches.

As Equation (14) shows, the smaller *m* is, the smaller the change rate becomes as the step-length increases. This means that when *m* is small, the occurrence of a long step-length becomes more frequent.The significant negative correlation result between the shape parameters of GE and the average autocorrelation coefficients was consistent with that of Wang et al. [23]. They found time dependence in the time-series data of step-lengths when its frequency distribution followed a power-law distribution. However, it is unclear why the shape parameter is associated with the time dependence. The shape parameter of the distribution considers the history of the distance traveled in a straight line, that is, how long the same process has lasted when elongating the straight-line distance, and is related to the process within single straight-line behavior. Conversely, the time dependence of the time-series data of straight-line distance is associated with the relationship between multiple straight-line behaviors. In the future, why this correlation is observed between the two, must be clarified theoretically and experimentally. It is also unclear why this association was observed in pill bugs and may have a completely different cause than in humans. The question of whether this relationship also manifests in the migratory behavior of animals other than pill bugs requires further research. The time dependence on step-length may occur when the shape and the exponent parameters are not fixed but vary with time or step-length.

The results shown in Figure 7 suggest that the relationship *β* ≈ −*m* + 1 approximately exists between the shape parameter *m* and the exponent *β* for some pill bugs. If this relationship holds, the GE represents the power-law distribution with exponent two in the case of *m* → 0 . This result is consistent with the LFFH, which states that search efficiency is maximized in the case of the Lévy walk with exponent two. However, as shown in Figure 3, the shape parameter *m* and the exponent *β* are theoretically independent in the GE. Thus, it is unclear why a relationship such as *β* ≈ −*m* + 1 is established in some pill bugs. We would like to address this issue in the future.

The GE handles the intermediate distribution between the exponential and power-law distributions. We defined the change rate *λ* (*l*) = *β l*^*m*−1^ as shown in Equation (14) to connect the exponential and power-law distributions. However, the change rate has countless definitions. Therefore, it is necessary to verify the validity and suitability of this definition in future work.

## Acknowledgments

We would like to thank Editage (www.editage.com) for English language editing.

## Author Contributions

**SS:** Conceptualization, formal analysis, methodology, software, writing—original draft preparation, funding acquisition.

**HO:** Conceptualization, methodology, writing—review, and editing.

**TM:** Resources, data curation, writing—original draft preparation.

**YN:** Writing—review and editing.

**TS:** Resources, data curation, writing—review, and editing.

**AU:** Resources, data curation, writing—review, and editing.

**UC:** Writing—review and editing, supervision, and project administration.

## Data Availability

The source code used to produce the results and analyses presented in this manuscript is available from the Dryad repository: https://datadryad.org/stash/share/v9hOyNs88WHHtOQDoZQ8GZ2MezBTsdMSuhGp2ZmYIHo For the pill bugs’ walking data, see the following reference: “Shokaku T, Moriyama T, Murakami H, Shinohara S, Manome N, Morioka K. Development of an automatic turntable-type multiple T-maze device and observation of pill bug behavior. Rev Sci Instrum. 2020;91:104104. DOI: 10.1063/5.0009531.”

